# A competence-regulated toxin-antitoxin system in *Haemophilus influenzae*

**DOI:** 10.1101/633164

**Authors:** Hailey Findlay Black, Scott Mastromatteo, Sunita Sinha, Rachel L. Ehrlich, Corey Nislow, Joshua Chang Mell, Rosemary J. Redfield

**Affiliations:** Department of Zoology, University of British Columbia, Vancouver BC, Canada; Sequencing + Bioinformatics Consortium, Office of the Vice-President, University of British Columbia, Vancouver BC, Canada; Department of Microbiology & Immunology, Center for Genomic Sciences, Drexel University College of Medicine, Philadelphia PA, USA; Department of Pharmaceutical Sciences, University of British Columbia, Vancouver BC, Canada

## Abstract

Natural competence allows bacteria to respond to environmental and nutritional cues by taking up free DNA from their surroundings, thus gaining both nutrients and genetic information. In the Gram-negative bacterium *Haemophilus influenzae*, the genes needed for DNA uptake are induced by the CRP and *Sxy* transcription factors in response to lack of preferred carbon sources and nucleotide precursors. Here we show that one of these genes, *HI0659*, encodes the antitoxin of a competence-regulated toxin-antitoxin operon (‘*toxTA’*), likely acquired by horizontal gene transfer from a *Streptococcus* species. Deletion of the putative toxin *(HI0660)* restores uptake to the antitoxin mutant. The full *toxTA* operon was present in only 17 of the 181 strains we examined; complete deletion was seen in 22 strains and deletions removing parts of the toxin gene in 142 others. In addition to the expected Sxy-and CRP-dependent-competence promoter, *HI0659/660* transcript analysis using RNA-seq identified an internal antitoxin-repressed promoter whose transcription starts within *toxT* and will yield nonfunctional protein. We propose that the most likely effect of unopposed toxin expression is non-specific cleavage of mRNAs and arrest or death of competent cells in the culture. Although the high frequency of *toxT* and *toxTA* deletions suggests that this competence-regulated toxin-antitoxin system may be mildly deleterious, it could also facilitate downregulation of protein synthesis and recycling of nucleotides under starvation conditions. Although our analyses were focused on the effects of *toxTA*, the RNA-seq dataset will be a useful resource for further investigations into competence regulation.

**ABBREVIATED SUMMARY:** The competence regulon of *Haemophilus influenzae* includes an unprecedented toxin/antitoxin gene pair. When not opposed by antitoxin, the toxin completely prevents DNA uptake but causes only very minor decreases in cell growth and competence gene expression. The TA gene pair was acquired by horizontal gene transfer, and the toxin gene has undergone repeated deletions in other strains.

## INTRODUCTION

Bacterial toxin-antitoxin gene pairs were originally discovered on plasmids, where they function to promote plasmid persistence by killing any daughter cells that have not inherited the plasmid. Typically, one gene of the pair encodes a relatively stable toxic protein that blocks cell growth, and the other encodes a labile antitoxin (RNA or protein) that blocks the toxin’s activity and limits its expression [1,2]. Toxin-antitoxin gene pairs have also been discovered on many bacterial chromosomes, where they are thought to be relatively recent introductions that in some cases have been co-opted to regulate cellular functions or provide other benefits [3]. Here we describe one such system, which is induced in naturally competent cells and whose unopposed toxin completely prevents DNA uptake and transformation.

Many bacteria are naturally competent, able to take up DNA from their surroundings and—when sequence similarity allows—recombine it into their genomes [4,5,6]. In most species, DNA uptake is tightly controlled, with protein machinery specified by a set of co-regulated chromosomal genes induced in response to diverse cellular signals. Genes in the competence regulon encode not only components of the DNA uptake machinery that moves DNA across the outer membrane of the cell, but proteins that translocate DNA across the inner membrane, proteins that facilitate recombination, and proteins of unknown function. *Haemophilus influenzae* has an unusually small and well-defined competence regulon (26 genes in 13 operons) induced by signals of energy and nucleotide scarcity [7,8]. Induction of these genes begins in response to depletion of phosphotransferase sugars. The resulting rise in cyclic AMP (cAMP) activates the transcription factor CRP, and the CRP/cAMP complex then stimulates transcription of genes with canonical CRP-promoter elements (CRP-N genes). Most of these genes help the cell to use alternative carbon sources, but one encodes the competence-specific transcriptional activator Sxy. However, efficient translation of *sxy* mRNA occurs only when purine pools are also depleted [9,10]. If both signals are active, Sxy then acts with CRP at the promoters of competence genes, stimulating their expression and leading to DNA uptake and natural transformation. These competence promoters are distinguished by the presence of ‘CRP-S’ sites (formerly called CRE sites), variants of standard CRP sites that depend on both CRP and Sxy for activation [11], but the specific role of Sxy is not yet understood. Development of competence thus requires the CRP/cAMP complex twice, first for *sxy* transcription (at its CRP-N promoter) and then for transcription of the competence genes (at their CRP-S promoters).

All but one of the fifteen Sxy-regulated *H. influenzae* genes needed for DNA uptake encode typical competence proteins — membrane-associated proteins homologous to known components of the Type IV pilus-based DNA uptake machinery present in nearly all known naturally competent species [5]. The one exception is *HI0659*, which instead encodes a predicted 98 amino acid cytoplasmic protein with no similarity to known DNA uptake proteins. It shares a competence-inducible CRP-S promoter with an upstream gene encoding another short cytoplasmic protein (*HI0660*, 119 aa) (**Fig. 1, top**). Although a knockout of *HI0659* eliminates detectable DNA uptake and transformation, a knockout of *HI0660* has no effect [8].

**Figure 1:**
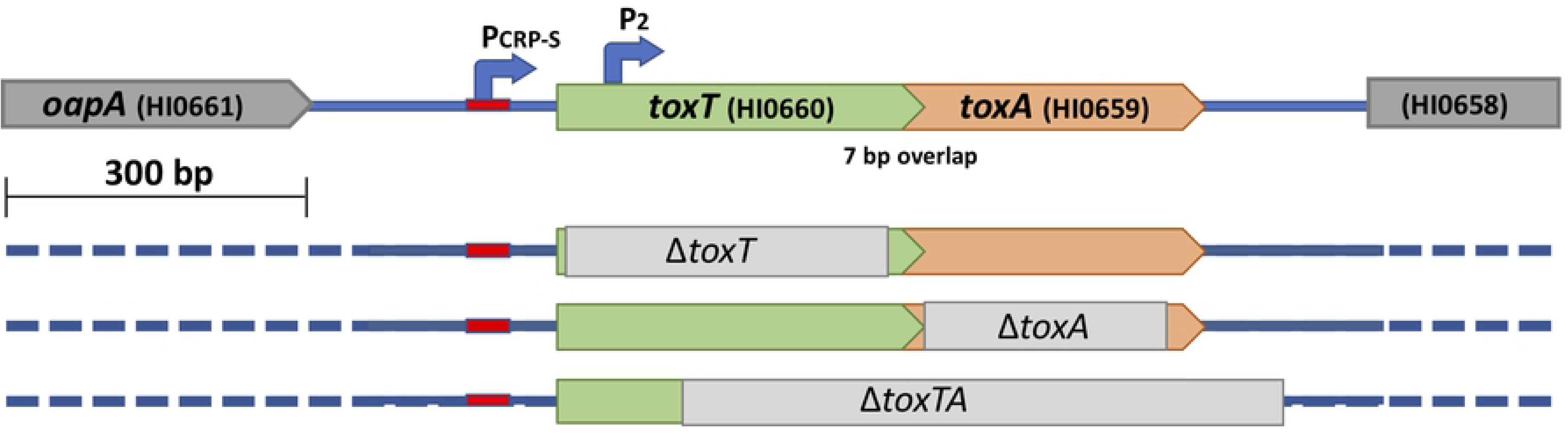
Structure of wildtype and mutant *toxTA* genes. Top line: structure of the wildtype *toxTA* operon in strain Rd KW20. Lower lines: grey bars indicate segments deleted in Δ*toxT*, Δ*toxA*, and Δ*toxT*A mutants.

Here we show that *HI0660* and *HI0659* comprise a horizontally transferred operon that encodes a toxin-antitoxin pair, and that expression of the toxin in the absence of the antitoxin completely prevents DNA uptake and transformation. Surprisingly, expression of this toxin has only modest effects on induction of competence genes and on cell growth and viability.

## RESULTS

### HI0659 and HI0660 act as a toxin-antitoxin system that affects natural competence

Our original analyses of competence-induced genes did not identify any close homologs of *HI0659* or *HI0660* [7,8]. However subsequent recent database searches and examination of BLASTP results revealed that these genes’ products resemble proteins in the Type II toxin/antitoxin families, which typically occur in similar two-gene operons. If *HI0660* and *HI0659* do encode a toxin-antitoxin pair, then *ΔHI0659*’s DNA uptake defect would likely be caused by unopposed expression of a *HI0660*-encoded toxin protein that prevents DNA uptake, and knocking out this toxin gene would restore competence to the *HI0659* (antitoxin^−^) mutant. We tested this by constructing an *HI0660/HI0659* double mutant (**Fig. 1**) and examining its ability to be transformed with antibiotic-resistant chromosomal DNA (MAP7) compared to wild type or either single mutant. The double mutant had normal transformation (**Fig. 2, black bars**), showing that mutation of *HI0660* suppresses the competence defect of an *HI0659* mutant, and also that neither *HI0660* nor *HI0659* is directly needed for the development of competence. This supported the postulated antitoxin function of *HI0660*, so we named the *HI0660* and *HI0659* genes *toxT* (toxin) and *toxA* (antitoxin) respectively.

**Figure 2:**
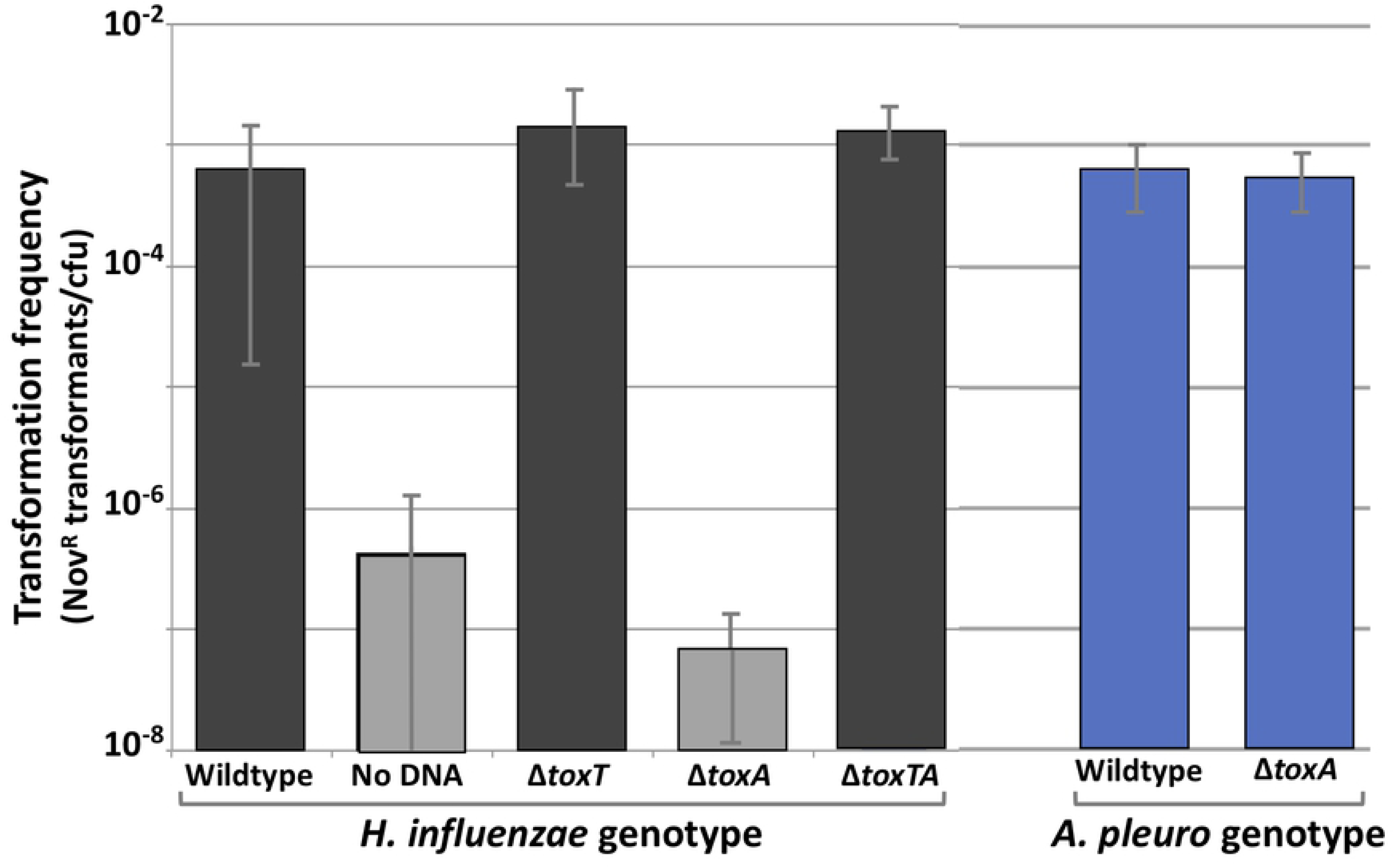
Transformation phenotypes of wildtype cells and *toxTA* mutants. MIV-induced cultures were incubated with MAP7 chromosomal DNA, and transformation frequencies were calculated as the novobiocin resistant (Nov^R^) colonies per colony forming unit (CFU). Bars represent the means of at least three biological replicates, with error bars representing one standard deviation. Grey bars indicate values below the detection limit (∼10^−8^ CFU/ml).

### Phylogenetic evidence for lateral transfer of *toxTA*

Since toxin/antitoxin operons are often highly mobile [13], we examined the distribution of *toxTA* homologs in other strains and species (Fig. 3). Complete or partial *toxTA* operons were found at the same genomic location in most *H. influenzae* genomes and in the closely related *H. haemolyticus* (see below), but there were no recognizable homologs in most other bacteria, including most other members of the Pasteurellaceae. Instead, most identifiable homologs (with about 60% amino acid identity) were in a very distant group, the Firmicutes, mainly within the streptococci. Ninety-six of the top 100 BLASTP hits to ToxT outside the Pasteurellaceae were to diverse *Streptococcus* species. This suggests that the *toxTA* operon may have been transferred from a Firmicute into a recent ancestor of *H. influenzae* and *H. haemolyticus*. When we excluded *Streptococcus* from the BLAST search, sporadic matches were found in a wide variety of other taxa. In addition, *toxTA* operons with about 50% identity were found in one other small distant Pasteurellacean subclade (*Actinobacillus sensu stricto*), and on two 11kb plasmids (pRGRH1858 and pRGFK1025) from an uncultured member of a rat gut microbiome and an uncultivated *Selenomonas* sp. The genes flanking the *A. pleuropneumoniae* homologs are unrelated to those flanking HI0659 and HI0660. The distribution of *toxTA* homologs across taxa is summarized in Fig. 3A.

**Figure 3:**
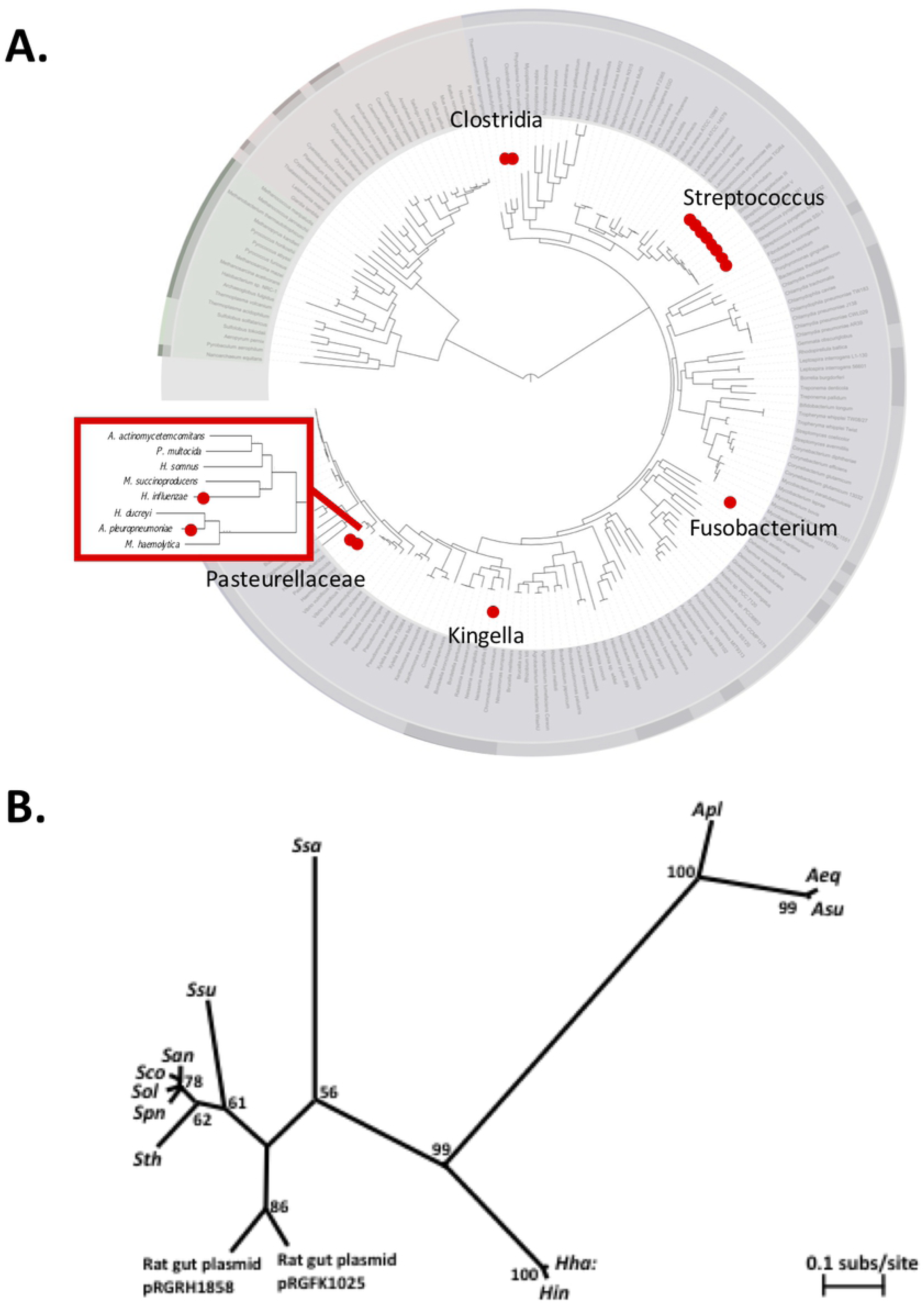
Distribution of *toxTA* homologs in bacterial genomes. **A.** Summary of BLASTP results. Red dots indicate one or more taxa containing homologs of both ToxT and ToxA. Bacterial phylogeny image from Wikimedia Commons [14]. Inset: Pasteurellacean phylogeny from Redfield *et al*. 2006. **B.** Unrooted maximum likelihood phylogeny of concatenated ToxT and ToxA homologs from selected species where both were detected. Numbers at nodes are bootstrap values. Species abbreviations: *Apl: Actinobacillus pleuropneumoniae; Aeq: A. equuli; Asu: A. suis; Haemophilus haemolyticus; Hin: H. influenzae; Ssa: Streptococcus salivarius; Ssu: S. suis; San: S. anginosus; Sco: S. constellatus; Sol: S. oligofermentans; Spn: S. pneumoniae; Sth: S. thermophilus.*

To resolve the history of gene transfer events in the two Pasteurellaceae sub-clades, we created an unrooted maximum likelihood phylogeny from the concatenated alignment of *toxT* and *toxA* amino acid sequences from selected species where both genes were found (Fig. 3B). Although a *Haemophilus*-*Actinobacillus* clade has 99% bootstrap support, the absence of homologs from all other Pasteurellaceae makes a single Pasteurellacean origin unlikely, since it would require multiple deletions in other Pasteurellacean subclades or a near-simultaneous within-clade lateral transfer. Since the *Actinobacillus* sequences are also more similar to *Streptococcus* sequences than to the *Haemophilus* sequences, the two Pasteurallacean groups are more likely to have acquired their *toxTA* operons by independent lateral transfers, from Firmicutes.

### Deletions in *H. influenzae toxT* are common

Of the 181 *H. influenzae* genome sequences available in public databases, 162 had recognizable *toxA* sequences. All of these encoded full length ToxA proteins, but all except 24 had one of two common deletions affecting *toxT*. The extents of these deletions are shown by the grey bars at the bottom of Fig. 4. The most common deletion (n=97) removed 178 bp of *toxT* coding sequence but left both promoters intact. The second (n=45) removed 306 bp of sequence including both *toxTA* promoters and the *toxT* start codon. The 19 genomes that lacked recognizable *toxA* sequences all had the same 1015 bp deletion removing both *toxT* and *toxA* but leaving the flanking genes intact. In place of the missing sequences were 87 bp with no high-scoring BLAST alignments in GenBank. The average pairwise distance among the 162 *toxA* genes is 0.106, slightly higher than the average of all genes with one copy per strain (0.088). The d_N_/d_S_ ratio of 0.037 is consistent with mild purifying selection on *toxA* and is higher than that of the average gene (0.243). However, this may underestimate the strength of selection, since most *toxA*s lack functional *toxT*, no strains with *toxT* only were seen, and the operon may not be expressed due to upstream deletions. Both sequence divergence and the high frequency of *toxT* deletions agree with expectations for a toxin/antitoxin system whose antitoxin protects against a toxin that is at least mildly deleterious.

**Figure 4:**
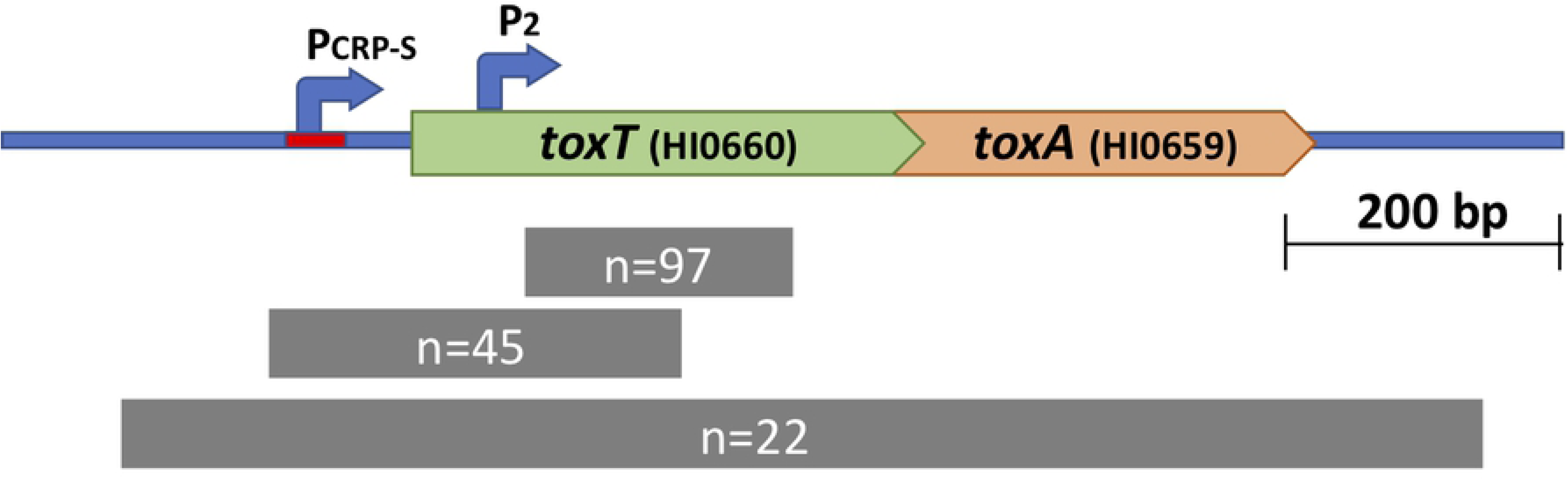
Natural deletions in the *toxTA* operon. Top line: structure of the wildtype *toxTA* operon in strain KW20. Lower lines: grey bars indicate the spans of the three naturally occurring deletions among the 181 available sequences, annotated with the number of sequences with each deletion

### Variation in *toxTA* is independent of strain-specific variations in DNA uptake and transformation

Maughan and Redfield [15] measured the ability of 34 *H. influenzae* strains to both take up DNA and become transformed, so we examined this data for correlations with the presence of *toxTA* in the 19 of these strains with genome assemblies whose *toxTA* genotypes we were able to determine. All but one of the 19 strains had a complete *toxA* coding sequence but only five had intact *toxTA* operons. Of the rest, four had the large deletion that removed both *toxTA* promoters, nine had the smaller deletion internal to *toxT*, and one had the 1015 bp complete deletion. There was no obvious correlation between the *toxTA* genotypes and the DNA uptake or transformation phenotypes, but there was insufficient data for a high-powered analysis using allelic variation.

### The *Actinobacillus pleuropneumoniae toxTA* operon does not control growth or competence

The *A. pleuropneumoniae toxTA* operon was originally reported to have the CRP-S promoter typical of competence operons [16]. Although reexamination of the promoter region failed to identify a high-quality CRP-S site, we constructed *toxT, toxA* and *toxTA* knockout mutants to investigate whether a *toxA* deletion would prevent competence. There were no significant differences between the transformation frequencies of wildtype cells and *toxA* mutants (Fig. 2, blue bars), and the rich-medium growth properties of all mutants in the wells of a BioScreen incubator were identical to those of wildtype cells (Supp. Fig. A). Thus, we conclude that the *toxTA* homolog in *A. pleuropneumoniae* does not affect competence. A previously reported RNA-seq study [17] of *A. pleuropneumoniae* serotype JL03 [18] found low-level expression of the *toxTA* homologs in a log-phase culture in rich medium, but competent cultures have not been examined.

### Growth and competence phenotypes of *H. influenzae toxTA* mutants

Earlier investigation of DNA uptake and transformation by the *H. influenzae toxA* knockout strain found that both were below the limit of detection (>100-fold reduction and >10^6^-fold reduction respectively) after the standard competence-inducing treatment [8] (see also **Fig. 2**), although there was no apparent growth defect in rich medium. Supp. Fig B confirms the non-transformability of Δ*toxA* cells both in log phase growth and at intermediate time points during competence induction.

A simple explanation for this defect would be that unopposed ToxT prevents competence, when not opposed by ToxA, by killing or otherwise inactivating the (competence-induced) cells in which it is expressed. To detect effects of unopposed expression of *toxT* toxin, we analyzed growth rates of wildtype and Δ*toxA* strains (Fig. 5) before, during, and after transfer to the competence-inducing starvation medium MIV. The first 60 min of Fig. 5 show that unopposed expression of *toxT* toxin slightly slows exponential growth in rich medium. A more detailed analysis of growth of rich-medium cultures using BioScreen wells is provided in Supp. Fig. C; the growth defect of the Δ*toxA* strain is barely detectable under this condition. This lack of a severe growth defect is not surprising; because the *toxTA* promoter is regulated by a CRP-S site, its expression (and thus ToxT production) might be limited to competent cells even in the absence of ToxA [7]. The grey-shaded portion of Fig. 5 shows cell growth after transfer to MIV (samples between 70 min and 165 min). Cells transferred to MIV usually undergo only one or two doublings, and deletion of *toxA* delays this but does not eliminate it (see also Supp. Fig. D). Cells returned to rich medium from MIV might be delayed in their recovery, if unopposed toxin expression during competence development kills cells or halts growth. However, both strains had similar recovery kinetics after a fraction of their culture was returned to rich medium, although Δ*toxA* cells again grew slightly slower than wildtype (175-230 min in Fig. 5). Supp. Fig. E shows the OD_600_ readings corresponding to the CFU/ml data shown in Fig. 5. These agree well, showing that differences in cell growth can be explained by the differences in cell division.

**Figure 5:**
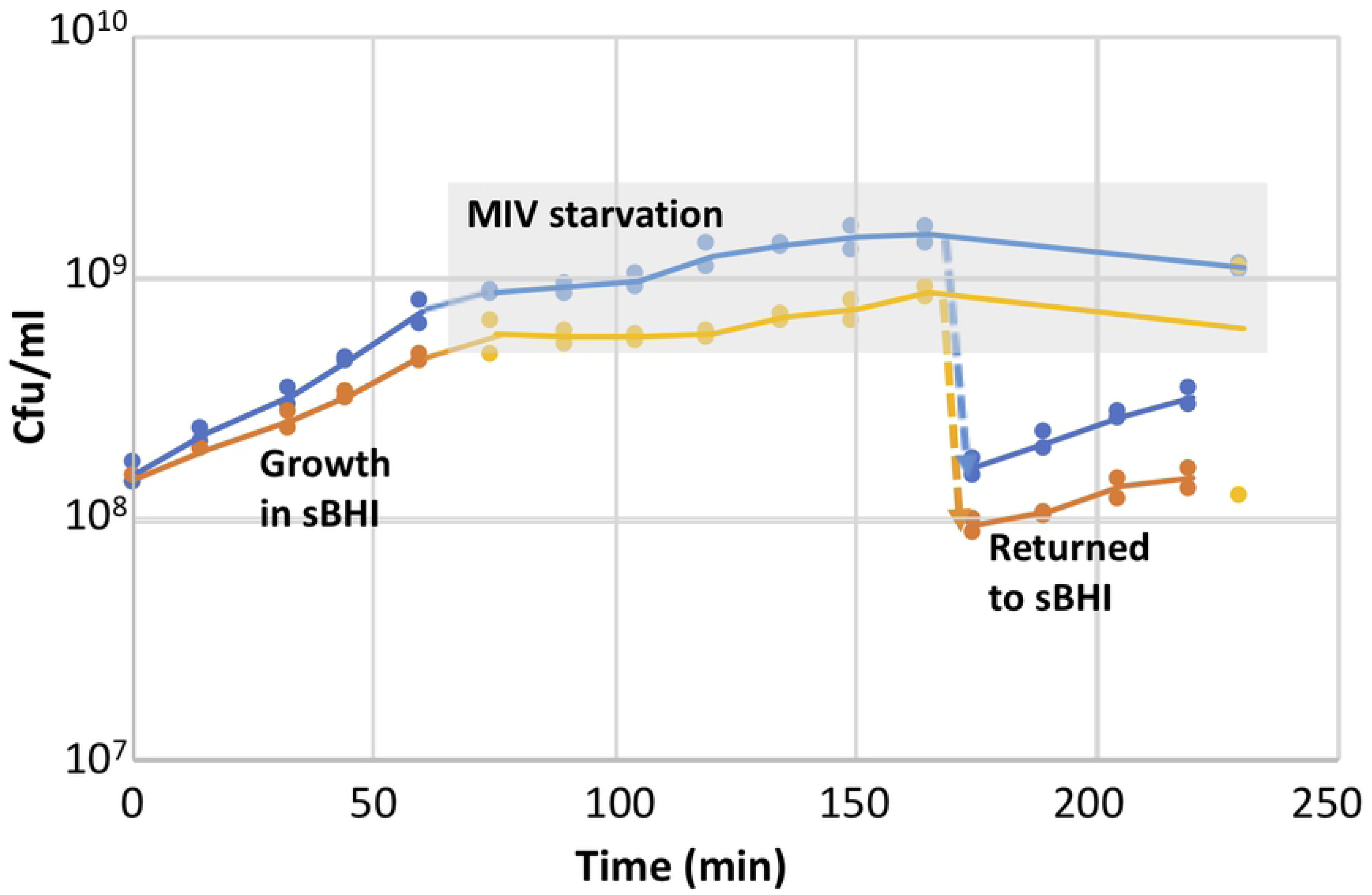
Growth of *H. influenzae* wildtype and Δ*toxA* cells before, during and after competence induction. Log-phase cells in sBHI were transferred to MIV at t=65 min; a portion of each MIV culture was diluted 10-fold into sBHI at t=170 min. The grey-shaded area indicates samples taken from MIV cultures. Blue: KW20, orange: Δ*toxA*. Supp. Fig. E shows the corresponding OD600 values.

Since cyclic AMP is required for normal induction of the competence genes, and addition of cAMP induces partial competence during exponential growth [19], we also tested the effect of cAMP on the Δ*toxA* knockout. Addition of cAMP did not rescue its transformation defect (Supp. Fig. F), so failure to transform is not caused by defective cAMP production in the antitoxin mutant.

Since some chromosomal toxin-antitoxin systems have acquired beneficial roles in modulating cell growth [20], we also examined whether the absence of toxin changed growth and competence under various conditions. The grey line in Supp. Fig. C shows that, under BioScreen growth assay conditions, Δ*toxT’s* growth was indistinguishable from that of wildtype cells (blue line), and Fig. 2 shows that its MIV-induced competence is also unchanged. Supp. Fig. G shows that the kinetics of Δ*toxT* competence development and loss during growth in rich medium were also indistinguishable from wildtype. We conclude that ToxT’s normal expression in cells expressing antitoxin does not detectably regulate growth or the development or loss of competence. Unfortunately, inferring the relationship between competence and growth is complicated because many cells in ‘competent’ cultures are unable to transform. In some species the non-transformable cells are known to have not induced their competence genes, but this has not been investigated for *H. influenzae*.

### Transcriptional control of competence

Since these phenotypic analyses gave little evidence of MIV-specific toxicity or insight into the cause of the competence defect, we used RNA-seq to investigate how *toxTA* is regulated and how mutations affect expression of competence genes. In these experiments, samples for RNA preparations were taken from three replicate cultures at four time points, first when cells were growing in log phase in the rich medium sBHI (t=0), and then at 10, 30 and 100 minutes after each culture had been transferred to MIV. We first examined how competence induction in wildtype and regulatory-mutant cells changed expression of genes known to be regulated by CRP and CRP+Sxy (CRP-N and CRP-S genes respectively). **Fig. 6** gives an overview of the results. The top row shows competence-induced changes in transcript abundances in wildtpe cells at 10, 30 and 100 minutes, and the lower rows show that some of these changes do not occur in Δ*crp* and Δ*sxy* cells. Each coloured dot represents a gene, colour-coded by function. Its horizontal position indicates its level of expression in rich medium (T=0) and its vertical position indicates how this expression changed in MIV (top row – wildtype cells, lower rows, Δ*crp* and Δ*sxy* cells). Thus in **Fig. 6A** the higher positions of the dark blue dots (genes regulated by CRP-N sites) and the red diamond (the competence regulator *sxy*) indicate that they were strongly induced after 10 min in MIV. Induction of *sxy* was followed at 30 and 100 minutes by strong induction of the known competence-regulon genes (higher positions of CRP-S genes; light green dots) (Fig. 6 B and C). Consistent with prior studies [7], induction of all these genes was blocked by deletion of the *crp* gene (**Fig. 6 D-F**), and induction of the competence regulon (CRP-S) genes was blocked by deletion of *sxy* (**Fig. 6 G-I**). Analysis of other competence-associated changes in gene expression in normal cells and in Δ*crp* and Δ*sxy* mutants is provided in the Supplementary Materials.

**Figure 6:**
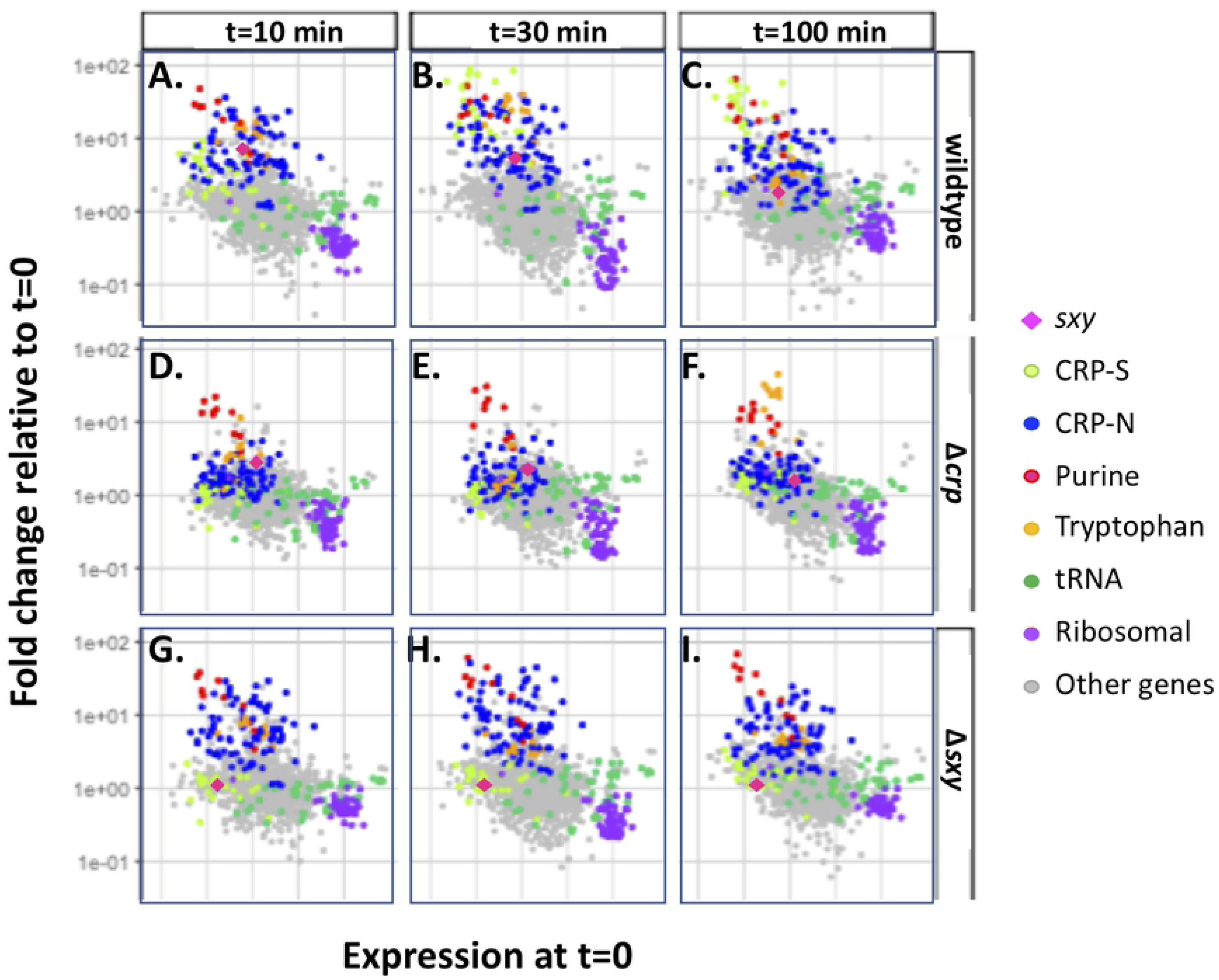
Competence-induced changes in expression of *H. influenzae* genes regulated by CRP and Sxy. Each dot represents a gene, colour-coded by function: pink diamond: *sxy*; light green dot: CRP-S-regulated; blue dot: CRP-N-regulated; red dot: purR regulon; yellow dot: tryptophan regulon; dark green dot: tRNA; purple dot: ribosomal proteins; grey dot: other or unknown functions. Each dot’s horizontal position indicates the gene’s relative level of expression (as FPKM) in rich medium (t=0) and its vertical position indicates how this expression changed at later time points (**A**: t=10; **B**: t=30; **C**: t=100) or in a mutant background at T=30 (**D-F**; Δ*crp*; **G-I**: Δ*sxy*).

### Transcriptional control of *toxTA*

RNA-seq analysis confirmed that *toxTA* is regulated as a typical competence operon. In wildtype cells, baseline RNA-seq expression of *toxT* and *toxA* was very low during log phase in rich medium, with approximately tenfold induction after 30 minutes incubation in MIV (green lines and points in **Fig. 7** (*toxT*) and Supp. Fig. H (*toxA*). As expected, this increase was eliminated by knockouts of CRP and Sxy (brown and blue lines and points in Fig. & and Supp. Fig H). Like other CRP-S genes, both *toxT* and *toxA* were also strongly induced in rich medium by *sxy* and *murE* mutations known to cause hypercompetence [38,39] and by a newly identified hypercompetence mutation in *rpoD* (**Supp. Fig. I**). Thus the *toxTA* operon is regulated as a typical member of the competence regulon.

**Figure 7:**
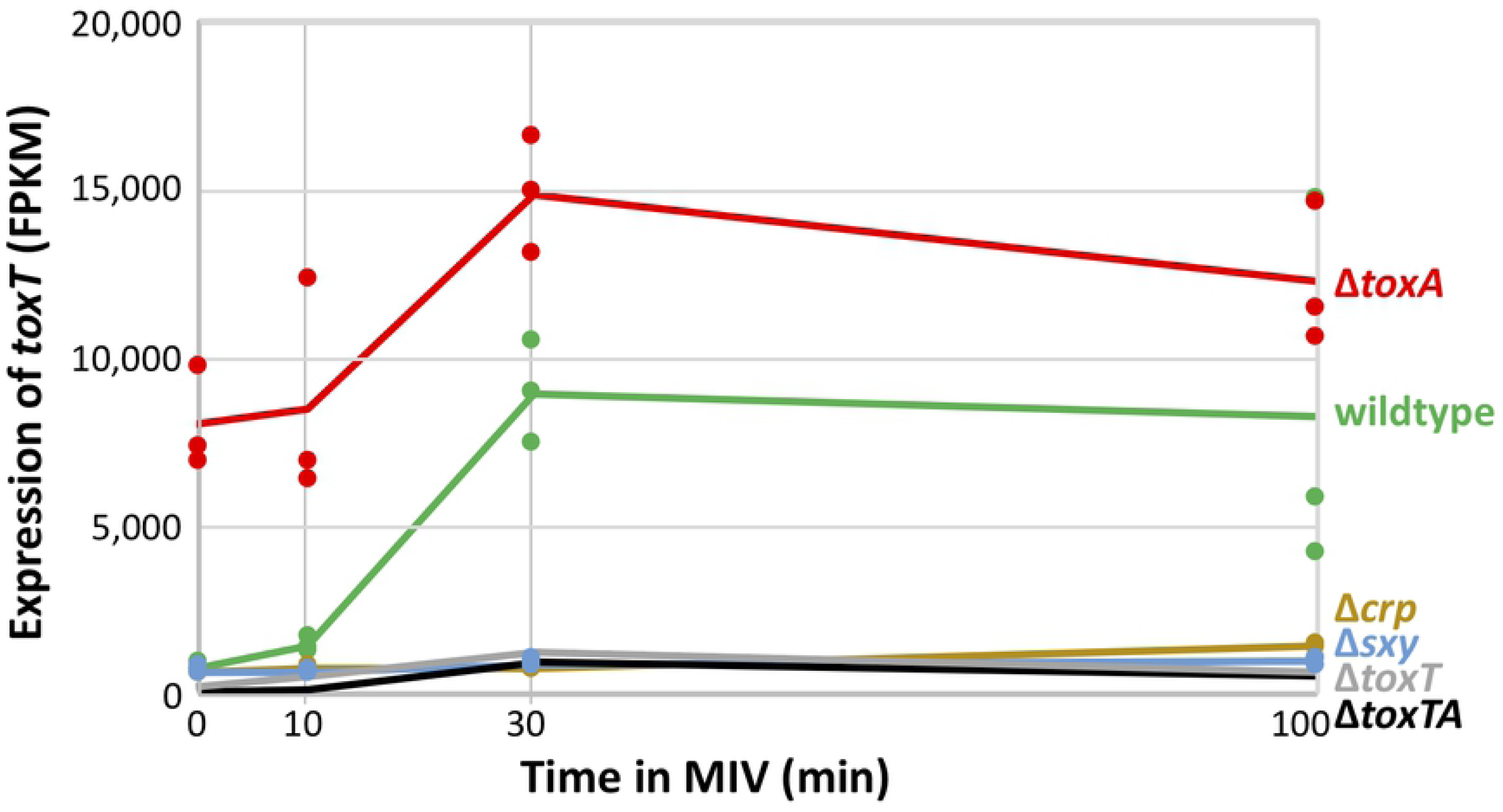
Competence-induced changes in expression of *H. influenzae toxT*. Sample FPKM values (dots) and means (lines) for *toxT* (HI0660). Strains: wildtype: green; Δ*crp*: brown; Δ*sxy*: blue; Δ*toxA*: red; Δ*toxT*: grey; Δ*toxTA*: black. The values for the Δ*toxT* and Δ*toxTA* samples are underestimates because most of the gene has been deleted in these strains.

RNA-seq analysis also showed that the *toxTA* operon is regulated as a typical type II toxin-antitoxin operon. In such operons, the antitoxin protein usually binds to the toxin protein, which protects cells from the toxin in two ways. First, antitoxin binding inactivates the toxin. Second, it also activates the antitoxin component as a repressor of the *toxTA* promoter [2,21]. *H. influenzae* ToxA has a HTH-XRE DNA-binding domain, which is commonly found in promoter-binding antitoxins [1,13], and the RNA-seq analysis in **Fig. 7** strongly suggests that it represses *toxTA* transcription. The Δ*toxA* mutant retains an intact *toxTA* promoter and *toxT* coding sequence (see **Fig. 1**); it had 9-fold increased baseline expression of *toxT* in log phase cells (red line and points in **Fig. 7**). Expression increased further during competence development, with the same kinetics as in wildtype cells, suggesting independent contributions from baseline repression by antitoxin and competence induction by CRP and Sxy. (Values for *toxA* expression are shown by the red points and line in **Supp Fig. H**, but are underestimates because most of the gene has been deleted.)

Because most antitoxins have only weak affinity for DNA in the absence of their cognate toxin, ToxA was predicted to repress *toxTA* only when bound to ToxT. Therefore, we were initially surprised that knocking out *toxT* or both *toxT* and *toxA* did not increase RNA-seq coverage of residual *toxT* sequences (**Fig. 7**, grey and black lines) and that knocking out both genes did not increase coverage of residual *toxA* (grey line in **Supp Fig. H**). These deletion mutants retain all the *toxTA* upstream sequences and the *toxT* start codon, and enough sequence of the deleted genes to identify them in the RNA-seq analysis. An explanation was suggested by a recent study of the *Escherichia coli hicAB* toxin/antitoxin system [22], and confirmed by more detailed analysis of *toxTA* transcripts. The HicA (toxin) and HicB (antitoxin) proteins have no detectable sequence homology to ToxT and ToxA, but their operon is similarly regulated by Sxy and has the same atypical organization (toxin before antitoxin) [23]. Turnbull and Gerdes [22] showed that the *hicAB* operon has two promoters. Promoter P1 has a CRP-S site regulated by CRP and Sxy, which is not repressed by the HicB antitoxin. A secondary promoter P2 is very close to the *hicA* start codon; it is repressed by HicB independently of HicA, and its shortened transcripts produce only functional HicB, not HicA. Promoter P1 of this *hicAB* system thus resembles the CRP-S regulation of the *toxTA* operon, and the presence of a second antitoxin-regulated internal promoter similar to P2 would explain the high *toxTA* operon expression seen in the *toxA* knockouts. This finding prompted us to do a more detailed analysis of *toxTA* transcription patterns in wildtype and mutant cells to determine whether the *toxTA* transcripts expressed in the absence of *toxA* were similarly truncated. **Figure 8A** shows RNA-seq coverage of the *toxTA* promoter region and the 5’ half of *toxT*, in wildtype cells (green) and in the *toxA* deletion mutant (purple). As expected, the predicted CRP-S promoter upstream of *toxTA* was only slightly active at T=10 but strongly induced at T=30 and T=100 (note log scale); its activity was not affected by deletion of *toxA*. Deletion of *toxA* instead caused strong constitutive transcription from a second promoter (‘P2’), with reads beginning about 30 bp downstream of the *toxT* start codon. Transcripts with this 5’ end are unlikely to produce active ToxT; the only other in-frame AUG in *toxT* is 30 bp from the end of the gene, and it and the first GUG (position 35) lack Shine-Dalgarno sequences. This supports the hypothesis that the *H. influenzae toxTA* operon is regulated similarly to the *E. coli hicAB* operon, with Sxy-induced expression from a CRP-S promoter and antitoxin-repressed transcription from a downstream ‘P2’ promoter whose transcript produces antitoxin but not toxin.

**Figure 8:**
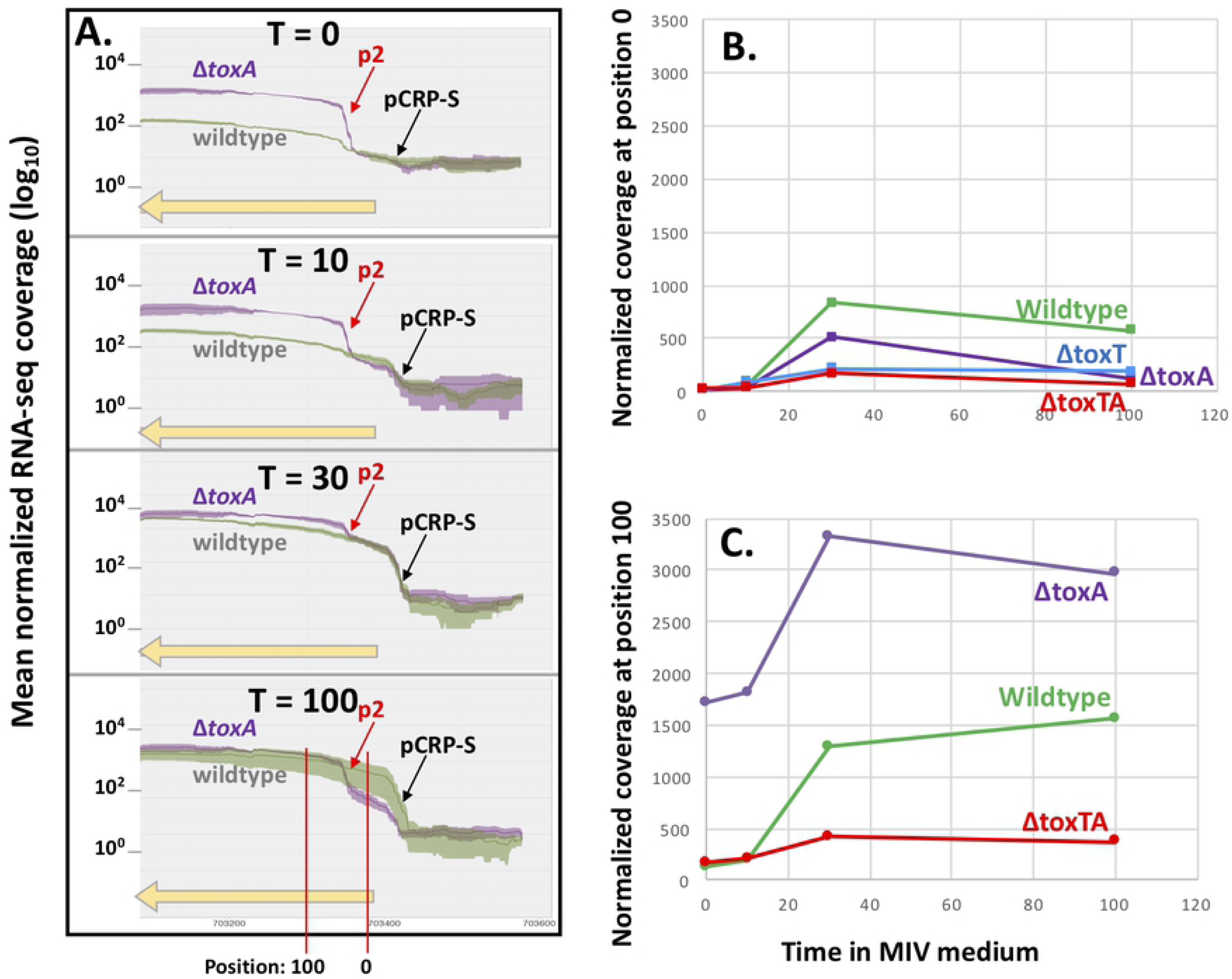
Sequencing read coverage of the *H. influenzae toxTA* promoter region. **A.** The green (KW20) and purple (Δ*toxA*) lines indicate mean coverage normalized by library size using DESeq2 [size factors] at each position; shaded areas indicate standard errors. The yellow horizontal arrow indicates the 5’ half of *toxT (*note that in this figure transcription is from right to left). **B.** and **C.** Time course of normalized read coverage at two specific positions in the *toxTA* operon. **B.** Position 0 = *toxA* start codon. **C.** Position 100.

In the *E. coli hicAB* system, P2 is repressed by HicB antitoxin alone, binding of HicB to the P2 operator is destabilized when HicA toxin is abundant, and transcription from P2 in plasmid constructs is elevated when the chromosomal *hicAB* operon is deleted [22]. To look for parallels in *H. influenzae*’s *toxTA*, we more precisely measured transcription in wildtype and *toxTA* mutant cells by scoring the coverage at two positions in the *toxTA* operon; each is indicated by a red vertical line at the bottom of Fig. 8A. Position 0 is the *toxT* start codon, 34 nt downstream from the CRP-S promoter (P_CRP-S_) but upstream of the putative P2 promoter, and position 100 is 70 nt downstream from P2 (P2 and position 100 are deleted in Δ*toxT*). To eliminate read-mapping artefacts arising from failure of reads that span an insertion or deletion to align to the reference sequence, each mutant’s reads were instead mapped onto its own *toxTA* sequence. Comparison of **Figures 8B and 8C** shows that coverage at position 100 was always higher than coverage at position 0, consistent with the presence of a second promoter between positions 0 and 100. **Fig. 8B** also shows that coverage at position 0 (expression from P_CRP-S_) was reduced by all of the *toxTA* deletions. This was unexpected, and suggests that this promoter may have unusual properties, since coverage of other CRP-S genes was not similarly affected. The *toxA* deletion caused the predicted increase in coverage at position 100 (**Fig. 8C**), but the *toxTA* deletion unexpectedly reduced rather than increased coverage at this position ∼3-fold from the wildtype level, even though this construct retains the first 150 bp of the operon, including P2. This reduction was not accounted for by the reduction in expression from P_CRP-S_, suggesting that high-level transcription from the *toxTA* P2 promoter only occurs when ToxT is present and ToxA is absent. This could mean either that ToxT directly binds the P2 promoter to induce transcription, which seems unlikely given its lack of DNA-binding domain, or that the presence of ToxT disrupts binding of a secondary repressor of the operon, such as a noncognate antitoxin [2]. Alternatively, it is possible that that the reduced expression in the Δ*toxTA* mutant instead reflects reduced transcript stability in this mutant.

### Unopposed ToxT does not block induction of the competence regulon

Expression of the competence operons that these regulators induce was also normal or near-normal in the Δ*toxA* mutant at 30 min after transfer to MIV, the time when competence-induced gene expression is normally highest (**Fig. 9**). Modest decreases in expression were seen for some operons, but these are not expected to cause the absolute competence defect, for two reasons. First, competence gene expression levels at this time were very similar between the Δ*toxA* mutant, which cannot take up any DNA or produce any transformants (dark blue bars), and the Δ*toxT* and Δ*toxTA* mutants (other blue bars), which have normal competence. Second, an unrelated regulatory defect mutation, a knockout of *hfq*, causes a more extreme reduction in gene expression at 30 min (brown bars) but this causes a much less extreme competence defect (only 10-20 fold rather than >10^6^-fold; green line in **Supp. Fig. B**). Mutation of *toxTA* genes also did not substantially change expression of the *sxy, crp* and *cya* genes encoding the competence regulon regulators Sxy, CRP, and adenylate cyclase **(Supp. Fig. J)**. At t=100 the Δ*toxA* mutant showed a stronger reduction in RNA-seq coverage of competence genes (**Supp. Fig. K**), however this cannot explain the competence defect and its significance is unclear.

**Fig. 9.**
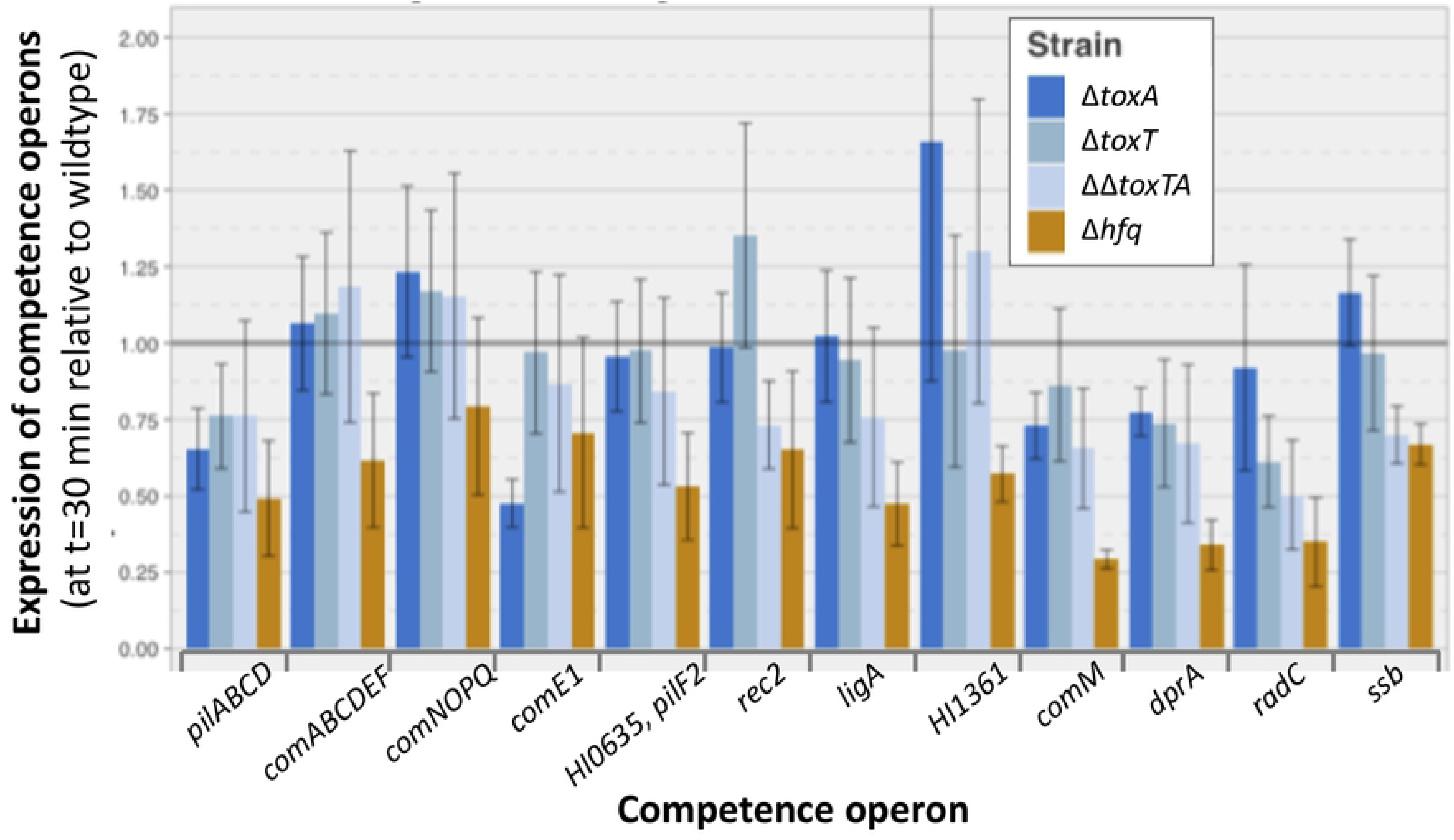
RNA-seq analysis of *H. influenzae toxTA* effects on competence-operon expression. Bars show the relative expression (as FPKM) of competence operons in four different mutants after 30 minutes of competence induction: Δ*toxA*: dark blue, Δ*toxT*: blue, Δ*toxTA*: light blue, Δ*hfq*: brown. Expression of each operon is relative to its expression in wildtype cells at the same time point (grey horizontal bar). Error bars indicate standard errors of samples from three replicate cultures.

### Other ToxT and ToxA effects in competence-induced cells

The severe competence defect of Δ*toxA* was also not explained by expression changes in other genes. Supp. Table 3 lists, for each timepoint, the genes whose expression in the Δ*toxA* mutant differed significantly from expression in all three strains with normal competence (wildtype, Δ*toxT* and Δ*toxTA*). The most consistent effect was overexpression of the HI0655-58 operon immediately downstream from *toxTA*. Analysis of sequencing coverage in this region showed this to be caused by read-through transcription from both *toxTA* promoters (CRP-S and P2). The operon includes genes encoding shikimate dehydrogenase, an ABC transporter, and a hypothetical protein with putative topoisomerase I domains. Expression of genes in this operon increased about 1.2-1.5-fold in MIV in wildtype cells and in other mutants with normal competence, so their higher expression in Δ*toxA* is unlikely to be responsible for this strain’s competence defect. Several other genes showed consistent changes relative to most or all competent strains, but these only sometimes reached statistical significance. HI0231 (*deaD*) encodes a DEADbox helicase involved in ribosome assembly and mRNA decay [24]; its expression in control strains fell rapidly on transfer to MIV, but levels in Δ*toxA* were about 50% higher at all time points. HI0235 contains an ArfA ribosome-rescue domain [25]; its expression was up 3-5-fold at t=30 and 2-3-fold at t=100 in the Δ*toxT* and Δ*toxTA* comparisons, but not in the KW20 comparison. HI0362 encodes a CRP-regulated iron-transport protein that normally increases in MIV but did not increase in *toxA* deletion mutants. HI0504 (*rbsB*, a ribose transporter component) is normally induced 20-fold more in MIV than other genes in its operon, but this increase was smaller (only 10-fold) in Δ*toxA*. Expression of HI0595 (*arcC*, carbamate kinase) normally falls 2-3-fold immediately after transfer to MIV, but the fall was greater in Δ*toxA*. 28 other genes were significantly changed only at t=100, but their expression patterns and predicted functions were diverse and did not suggest an explanation for Δ*toxA*’s lack of competence.

### Related toxins may suggest mechanism of action for ToxT

Since examination of gene expression shed little light on how the ToxT toxin prevents competence, and it has no close homologs with characterized functions, we considered the modes of action of other Type II toxins. The most common type II toxins (e.g. RelE) act as translation-blocking ribonucleases, but several alternative modes of action are also known, and some newly discovered toxins lack identified activities [13]. The Pfam and TAfinder databases assign the *H. influenzae* ToxT protein to the ParE/RelE toxin superfamily, whose characterized members include both gyrase inhibitors and ribonucleases that arrest cell growth by cleaving mRNAs and other RNAs [2,26,27].

### Unopposed toxin does not inhibit gyrase

If ToxT inhibited gyrase we would expect the RNA-seq data to show that transfer to MIV caused increased expression of *gyrA* (HI1264) and *gyrB* (HI0567) and reduced expression of *topA* (HI1365), since these genes have opposing activities and compensatory regulation by DNA supercoiling [28]. However, these genes’ coverage levels were similar in wildtype and all *toxTA* mutants, during both exponential growth and competence development.

### Unopposed toxin does not cleave competence-induced mRNAs at specific sites

The best-studied homologs of the *toxT* toxin act by cleaving mRNAs at positions near their 5’ ends during their translation on the ribosome [29,30]. Because the resulting ‘non-stop’ mRNAs lack in-frame stop codons and cannot undergo the normal ribosome-release process, this causes a general block to translation [31] which is predicted to arrest cell growth until normal translation can be restored [32]. Thus we considered whether ToxT might prevent competence by one of two mechanisms. First, ToxT might specifically cleave the 5’ ends of competence-gene transcripts, eliminating their function without significantly changing their overall RNA-seq coverage levels or otherwise interfering with essential cell functions. Visual examination of RNA-seq coverage of all positions within the competence operons did not reveal any anomalies that might indicate that the mRNA in Δ*toxA* cells had been inactivated either by cleavage at specific sites or by random cleavage near the 5’ end [33]. As an example, Supp Fig. L compares read coverage across the *comNOPQ* operon in wildtype and Δ*toxA* cultures after 30 min in MIV. To detect nonspecific cleavage we examined the insert sizes of our RNA-seq sequencing libraries by comparing the spanning-length distributions of paired-end sequencing reads among strains. Because independent library preparations had different insert sizes, comparisons were limited to samples prepared at the same time. Fig. 10 shows that the Δ*toxA* samples from library batch 1 had shorter fragment sizes than the KW20 samples from the same batch, and that the difference increased as the time after competence induction increased. This supports the hypothesis that the extreme lack of competence in Δ*toxA* cultures is due to non-specific ToxT cleavage of mRNAs. This mechanism is consistent with those of homologous HigB/RelE toxins, which inhibit translation by cleavage of mRNA at the ribosomal A-site [34,35].

**Fig. 10.**
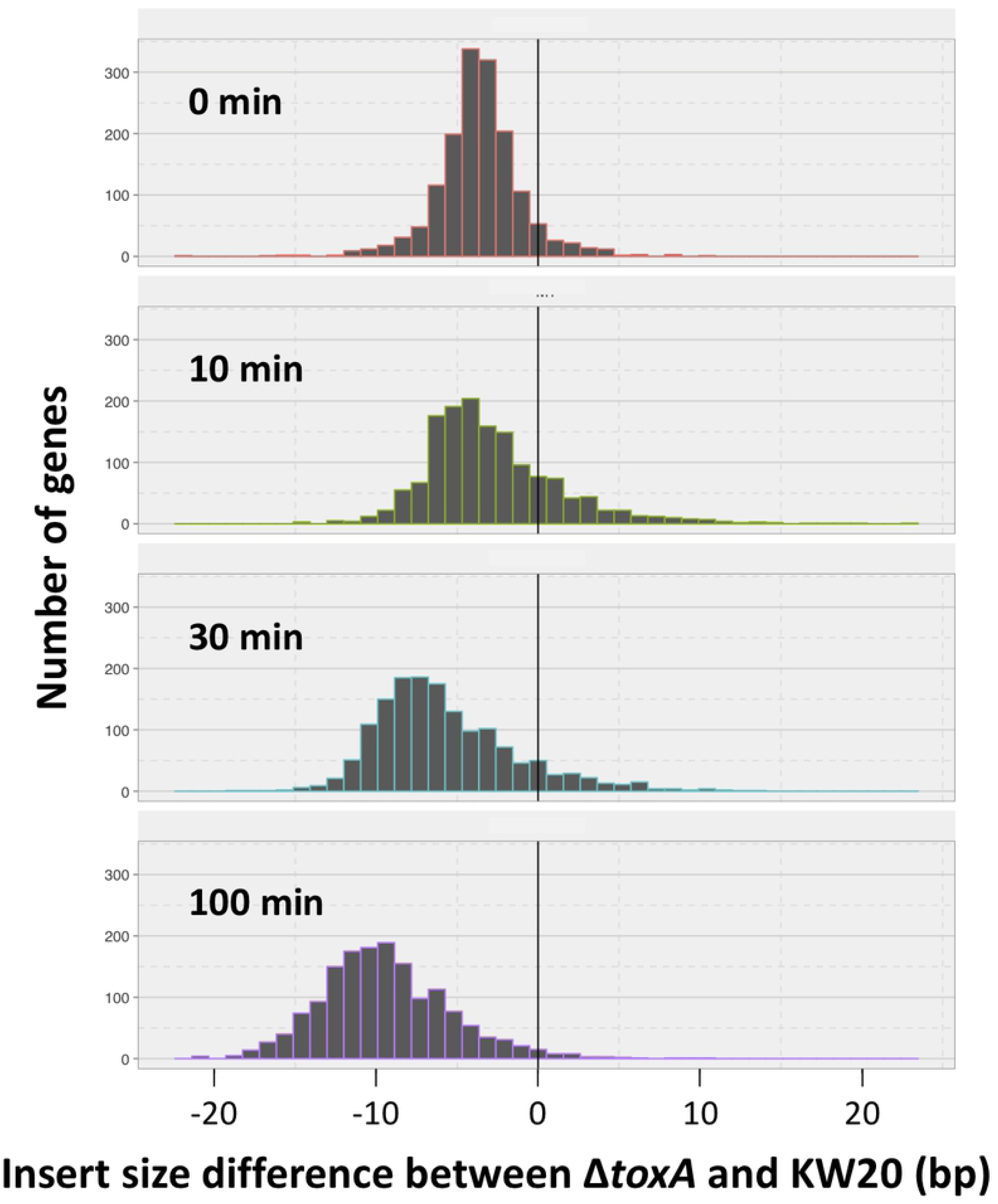
Distribution of insert-size differences between *H. influenzae* RNA-seq libraries prepared at the same time. Distribution of insert length differences between KW20 (kw20_A and kw20_B samples) and Δ*toxA* (antx_A samples) after 0, 10, 30 and 100 minutes in MIV.

## DISCUSSION

Our investigation into why a HI0659 knockout prevents competence has provided a simple answer: HI0659 encodes an antitoxin (ToxA) needed to block the expression and competence-preventing activity of the toxin encoded by HI0660 (ToxT). But this answer has generated a number of new questions that we have only been able to partially answer. Why is competence controlled by a toxin/antitoxin system? How does it completely abolish DNA uptake and transformation without causing significant cell death? Do its effects in wildtype cells confer any benefit to the cells, either generally or specific to competence?

Several findings support the conclusion that HI0660 and HI0659 encode proteins that function as a toxin/antitoxin pair. First is the similarity of the encoded ToxT and ToxA proteins to biochemically characterized toxin and antitoxin proteins of the RelE/ParE families. Second, and the strongest evidence, is the restoration of normal DNA uptake and transformation to antitoxin-knockout cells when the putative toxin is also knocked out. Third is the regulatory similarity between this system and the *hicAB* system of *E. coli*.

### How did the *toxTA* operon come to be in the *H. influenzae* genome and under competence regulation?

Phylogenetic analysis showed that *H. influenzae* acquired its *toxTA* operon by horizontal transfer, either into a deep ancestor of the Pasteurellaceae or more recently by independent transfers into ancestors of *H. influenzae* and *A. pleuropneumoniae*. The closest relatives of the *H. influenzae toxTA* genes are in the distantly related Firmicutes, with homologs especially common in *Streptococcus* species. Since the Streptococci and Pasteurellaceae share both natural competence and respiratory-tract niches in many mammals, there may have been frequent opportunities for horizontal transfer between them.

We do not know how the *toxTA* operon came to be under CRP-S regulation. The *toxTA* operon’s strong regulatory parallels with the *E. coli hicAB* system suggest that toxin-antitoxin systems with similar regulation and function have adopted similar roles in separate instances, a phenomenon which is more likely in toxin antitoxin systems as they undergo frequent horizontal transfers and are often under strong selective pressure. The *sxy* gene and the CRP-S promoters it regulates are not known outside of the Gamma-Proteobacteria sub-clade that contains the Vibrionaceae, Enterobacteraceae, Pasteurellaceae and Orbaceae [11]. Thus, it would be interesting to examine the regulation and function of the *toxTA* homologs outside the Pasteurellaceae to determine when and where it adopted a regulatory role and the mechanism of the toxic activity. Examining these homologs could give insight into both the mechanism of action of the *H. influenzae toxTA* system, and its evolutionary history.

### How does unopposed ToxT prevent DNA uptake and transformation?

The transformation defect caused by deletion of the antitoxin gene toxA is very severe, so it was surprising that RNA-seq analysis detected only few and minor changes in expression of competence genes. Instead, the best explanation is that ToxT is an mRNA-cleaving ribonuclease, whose activity causes a general block to translation that prevents functioning of the induced competence genes. The most direct evidence is the decrease in insert size distributions seen in Δ*toxA* mutants, but this conclusion is also supported by the combination of regulatory similarities between the *toxTA* and *hicAB* systems and by sequence similarities between the ToxT protein and HigB ribonuclease toxins.

### Why then does the Δ*toxA* mutant not suffer from growth arrest or toxicity?

Part of the explanation is that mRNAs encoding functional ToxT are only expressed after cells have been transferred to competence-inducing starvation medium, a condition that severely slows cell growth and division even in wildtype cells. Detecting the predicted competence-specific toxicity is further complicated by the uneven distribution of transformability in competence-induced cells. Co-transformation experiments using multiple unlinked markers consistently show that no more than half, and sometimes as little as 10%, of the cells in a MIV-treated culture produce recombinants [6]. We do not know whether only the transforming cells express the competence genes or all cells express them but some fail to correctly assemble the DNA uptake or recombination machinery. If only a modest fraction of the cells in a competent culture are expressing the toxin then any toxic effect on culture growth and survival will be more difficult to detect.

### Does this operon confer any benefit (or harm) on *H. influenzae*?

Why have a competence-regulating toxin/antitoxin system at all, when it has no detectable effect on competence unless its antitoxin component is defective? Although regulatory parallels with the *hicAB* system suggest that CRP-S regulation is not incidental, we found no direct evidence of any toxin-dependent alteration to the normal development of competence. Production of Sxy is subject to post-transcriptional regulation by the availability of nucleotide precursors [9,10], and we have elsewhere proposed that DNA uptake is an adaptation to obtain nucleotides when nucleotide scarcity threatens to arrest DNA replication forks [6]. In this context, competence-induction of the *toxTA* operon may be a specialization to help cells survive, by slowing or arresting protein synthesis until the nucleotide supply is restored.

On the other hand, the high frequency of deletions that remove either complete *toxTA* or both promoters (35%) indicates that the operon is dispensable. And the even higher frequency of toxin-inactivating deletions in the presence of intact antitoxin genes and CRP-S promoter (51%), coupled with the absence of any deletion that inactivates antitoxin but preserves toxin, indicates that unopposed toxin is indeed harmful under some natural circumstances.

We have examined the *toxTA* operon from many angles and answered our initial question of why *toxA* knockout prevents competence in *H. influenzae*, but have raised new questions whose eventual answers we hope will give us greater insight not just into the *toxTA* system, but competence regulation in general. A number of desirable follow-up experiments could improve understanding of the *toxTA* system, particularly complementation experiments to determine whether ToxA expression in Δ*toxA* can restore competence, and whether ToxT expression in Δ*toxTA* can block competence. In future work, it would also be valuable to express ToxT and ToxA in *E. coli* on separate plasmids to examine the system’s toxicity and effect on competence.

## METHODS

### Bacterial strains, plasmids, and growth conditions

Bacterial strains used in this work are listed in Supp Table 1. *Escherichia coli* strain DH5*α* [F80*lacZ* #(*lacIZYA-argF*) endA1] was used for all cloning steps; it was cultured in Luria-Bertani (LB) medium at 37°C and was made competent with rubidium chloride according to the method provided in the QIAexpressionist manual protocol 2 (Qiagen). When antibiotic selection was required, 100 µg/mL ampicillin and 50µg/mL spectinomycin were used.

*Haemophilus influenzae* cells were grown in sBHI medium (Brain Heart Infusion medium supplemented with 10mg/mL hemin and 2mg/mL NAD) at 37°C in a shaking water bath (liquid cultures) or incubator (plates). *H. influenzae* strain Rd KW20 [36], the standard laboratory strain, was used as the wild type for all experiments. Mutant strains used in this study were marked deletion mutants in which the coding region of the gene was replaced by a spectinomycin resistance cassette, as well as unmarked deletion mutants derived from these strains; the generation of these mutant strains is described in [8]. Specifically, we used an unmarked deletion of HI0659 (HI0659-), marked and unmarked deletions of HI0660 (HI0660::spec, HI0660-), and a marked deletion of the whole operon (HI0659/HI0660::spec). Knockout mutants of *crp* and *sxy* have been described previously [37,38].

*Actinobacillus pleuropneumoniae* cells were grown in BHI-N medium (Brain Heart Infusion medium supplemented with 100µg/mL NAD) at 37°C. *A. pleuropneumoniae* strain HS143 [40] was used as the wild type for all experiments. Marked deletion mutants in which the gene of interest was replaced by a spectinomycin resistance cassette strains were generated for this study as described below. The HS143 genome region containing the homologs of the *Actinobacillus pleuropneumoniae serovar 5b strain L20* APL_1357 and APL_1358 genes, plus approximately 1 kb of flanking sequence on each side, was PCR-amplified, ligated into Promega pGEM-T Easy and transformed into *E. coli*. Plasmid regions containing APL_1357, APL_1358, or both genes were deleted from the pGEM-based plasmid by inverse PCR, and the amplified fragments were blunt-end ligated to the spectinomycin resistance cassette [41] from genomic DNA of a *H. influenzae comN*::*spec* strain [8]. Plasmids linearized with ScaI were transformed into competent *A. pleuropneumoniae* HS143 and transformants were selected for spectinomycin resistance using 100µg/mL spectinomycin after 80 minutes of growth in nonselective medium.

### Generation of competent stocks

To induce competence, *H. influenzae* and *A. pleuropneumoniae* were cultured in sBHI or BHI-N respectively and transferred to competence-inducing medium MIV [42] when they reached an optical density at 600nm (OD_600_) of approximately 0.25 [43]. After incubation with gentle shaking at 37°C for a further 100 min (*H. influenzae*) or 150 min (*A. pleuropneumoniae*), cells were transformed or frozen in 16% glycerol at −80 °C for later use.

### Transformation assays

#### Transformation of MIV-competent cells

Transformation assays were carried out as described by Poje and Redfield [43]. MIV-competent *H. influenzae* or *A. pleuropneumoniae* cells were incubated at 37°C for 15 minutes with 1µg/ml DNA, then DNaseI (10µg/mL) was added and cultures were incubated for 5 minutes to ensure no DNA remained in the medium. *H. influenzae* cultures were transformed with MAP7 genomic DNA [44], which carries resistance genes for multiple antibiotics, while *A. pleuropneumoniae* cultures were transformed with genomic DNA from an *A. pleuropneumoniae* strain with spontaneous nalidixic acid resistance (generated in this lab). Cultures were diluted and plated on both plain and antibiotic-containing plates (2.5ug/mL novobiocin for *H. influenzae* cultures, 20ug/mL nalidixic acid for *A. pleuropneumoniae* cultures) and transformation frequencies were calculated as the ratio of transformed (antibiotic-resistant) cells to total cells. For *A. pleuropneumoniae*, transformed cells were given 80 minutes of expression time in BHI-N before plating.

#### Time courses in rich medium

*H. influenzae* cells from frozen stocks of overnight cultures were diluted in fresh sBHI and incubated with shaking at 37°C. Periodically, the OD_600_ was measured, and at predetermined optical densities aliquots of the culture were removed and transformed with MAP7 DNA and plated as described above.

#### Bioscreen Growth Analysis

The Bioscreen C apparatus (BioScreen Instruments Pvt. Ltd.) was used to measure growth. Cells frozen from overnight cultures were pre-grown at low density in sBHI, and 300µL aliquots of 100-fold dilutions were placed into 20 replicate wells of a 100-well Bioscreen plate. Wells at the edges of the plate were filled with medium alone as controls. Cells were grown in the Bioscreen at 37°C for 18 hours with gentle shaking, and OD_600_ readings were taken every 10 minutes. Readings were corrected by subtracting the OD_600_ measured for medium-only controls, and replicates for each strain were averaged at each time point to generate growth curves. Doubling times were calculated for each strain from the subset of time points that represents exponential growth phase, as determined by linearity on a semi-log plot of time versus OD_600_.

#### Competence growth and survival time course

Cells were grown in sBHI to a density of ∼2×10^8^ cfu/ml (OD600 = 0.075) and transferred to MIV. After 100 min (time for maximum competence development, an aliquot of each culture was diluted 1/10 into fresh sBHI for recovery and return to normal growth. A fraction of each culture was incubated in a shaking water bath, and aliquots of the initial and ‘recovery’ sBHI cultures were also grown and monitored in a Bioscreen incubator.

#### Cyclic AMP competence induction

*H. influenzae* cells in sBHI were incubated with shaking to an OD_600_ of approximately 0.05. Cultures were split and 1mM cAMP was added to one half. At an OD_600_ of approximately 0.3, aliquots were transformed with MAP7 DNA and plated as described above.

#### Phylogenetic Analysis

A nucleotide BLAST search (discontinuous MEGABLAST) and a protein BLAST search against translated nucleotide databases (tBLASTn) were used to identify homologs of the HI0659 and HI0660 genes [45]. Protein sequences found by the tBLASTn search were retained for analysis if they showed greater than 60% coverage and greater than 40% identity to the *H. influenzae* query sequence. For species with matching sequences in multiple strains, the sequence from only one strain was kept.

For species in which homologs of HI0659 and HI0660 were found next to one another, amino acid sequences of concatenated matrices were aligned by multiple-sequence alignment using MAFFT, version 7.220 [46], run from modules within Mesquite version 3.02 [47]. The L-INS-I alignment method was used due to its superior accuracy for small numbers of sequences. After inspection of the alignments, poorly-aligning sequences were removed from the analysis, and alignment was repeated.

Phylogenetic trees were generated using the RAxML [48] maximum likelihood tree inference program, run via the Zephyr package of Mesquite. For each gene, 50 search replicates were conducted, using the PROTGAMMAAUTO option to allow RAxML to automatically select the best protein evolution model to fit the data. Since these trees were found to correspond exactly to a set of trees generated using the PROTGAMMAJTT model, this faster model was used to generate a majority-rules consensus tree from 1000 bootstrap replicates for each gene.

#### Analysis of natural deletions

181 publicly available *H. influenzae* genomes were downloaded from NCBI and the Sanger centre. Genomes were re-annotated using Prokka v1.11 [49], and the pangenome was calculated using Roary v3.5.1 [50] with a minimum blastp threshold of 75. The *toxA* gene cluster in the pangenome was identified by finding the gene cluster that contained the *toxA* gene from Rd KW20, and the *hicA* cluster was identified by finding the gene cluster that contained the *hicA* gene from PittAA. 2300 bp genome sequences centered on *toxA* and/or *hicA* were extracted from all *H. influenzae* genomes containing recognizable *toxA* and/or *hicB* genes, and aligned by MAFFT. For strains that lacked recognizable *toxA* or *hicB*, sequences adjacent to the genes that normally flanked each operon were extracted. K_a_/K_s_ and pairwise distance were calculated for each gene using SeqinR v 3.4-5 [51] with codon aware gene alignments were made using Prank (v.100802).

### RNA-seq analysis

#### Sample Preparation

Cell cultures of *H. influenzae* strain Rd, Δ*crp* and Δ*sxy* derivatives, and Δ*toxTA* mutants were grown in sBHI to an OD_600_ of 0.2 – 0.25, then transferred to MIV. Aliquots of cells were removed just prior to transfer to MIV, and after 10, 30, and 100 minutes in MIV, and immediately mixed with Qiagen RNAprotect (#76526) to stabilize RNA. Cells were pelleted and frozen, and RNA was later extracted from thawed pellets using the Qiagen RNeasy Min-elute Cleanup Kit (#74204). Contaminating DNA was removed with Ambion Turbo DNase (#AM2238), and ribosomal RNA was depleted using the Illumina Ribo-Zero rRNA Removal kit (#MRZMB126). Sequencing libraries were prepared using TruSeq mRNA v2 library preparation kit, according to manufacturer’s instructions (Illumina). Libraries were pooled and sequenced on a HiSeq 2500, generating paired-end 100 bp reads.

#### Data Analysis Pipeline

FASTQ files were analysed using the FASTQC tool (Andrews, 2015) to confirm read quality. Reads were aligned to the *H. influenzae* Rd KW20 reference genome sequence using the Burrows-Wheeler Alignment tool (BWA) algorithm `bwa mem` [52]. Differential expression analysis was performed using the DESeq2 package, v.1.6.3 (Love *et al.*, 2013). Specifically, the function `DESeqDataSetFromMatrix()` was used to generate a dataset to compare reads from each mutant strain reads from the wild-type control based on their strain, sample time point, and the interaction between the two parameters. The function `DESeq()` was called to determine which genes were differentially expressed based on these parameters, using p-values adjusted for a B-H false-discovery rate [53] of 0.1 as a cut-off to determine significance, after normalizing total read counts and variances.

## AUTHOR CONTRIBUTIONS

- Performed the experiments: HFB, SM, SS, RJR
- Carried out the analyses: HFB, SM, SS, RLE, JCM, RJR
- Wrote the paper: HFB, SM, JCM, RJR
- Contributed reagents and supplies: CN, RJR

## Data availability

RNA-seq data were deposited under BioProject 293882

## ACKNOWLEDGEMENTS

We thank Lauri Lintott for helpful discussions, Charles Thompson for the use of the BioScreen, and Anni Zhang and Yvonne Yiu for technical assistance. This work was supported by funding from Canadian Institutes of Health Research to RJR, an NIH F32 AI084427 grant to JCM, and NIH R01 DC002148 to Garth D. Ehrlich. Sequencing work was performed at the Sequencing and Bioinformatics Consortium at the University of British Columbia.

## Supplementary Figure Captions

**Supplementary Figure A: Bioscreen analysis of growth of *A. pleuropneumoniae* wildtype and *toxTA* deletion strains in rich medium.**

**Supplementary Figure B. Development of competence by *H. influenzae* strains in MIV competence medium.**

**Supplementary Figure C: Bioscreen analysis of growth of *H. influenzae* wildtype and *toxTA* deletion strains in rich medium.**

**Supplementary Figure D. Growth of *H. influenzae* wildtype and Δ*toxA* strains after transfer to MIV competence medium.**

**Supplementary Figure E: Growth and MIV recovery of *H. influenzae* KW20 and ΔtoxA measured by OD**_**600**_.

**Supplementary Figure F: Effect of cAMP on *H. influenzae toxTA* mutants.**

**Supplementary Figure G: Transformation frequencies of wildtype and Δ*toxT* cells growing in rich medium.**

**Supplementary Figure H: Competence-induced expression of *H. influenzae toxA*.**

**Supplementary Figure I: Effect of hypercompetence mutations on expression of *H. influenzae toxA* and *toxT.***

**Supplementary Figure J: Effects of Δ*toxTA* mutations on expression of competence regulator genes.**

**Supplementary Figure K: Changes in expression levels of competence operons at t=100.**

**Supplementary Figure L: RNA-seq coverage of *H. influenzae comNOPQ* after 30 min of competence induction.**

